# Structure, substrate selectivity determinants and membrane interactions of a Glutamate-specific TAXI TRAP binding protein from *Vibrio cholerae*

**DOI:** 10.1101/2024.03.22.586268

**Authors:** Joseph F.S. Davies, Andrew Daab, Nicholas Massouh, Corey Kirkland, Bernadette Strongitharm, Andrew Leech, Marta Farré, Gavin H. Thomas, Christopher Mulligan

## Abstract

Tripartite ATP independent periplasmic (TRAP) transporters are widespread in prokaryotes and are responsible for the transport of a variety of different ligands, primarily organic acids. TRAP transporters are secondary active transporters that employ a substrate binding protein to bind and present the substrate to membrane embedded translocation component. TRAP transporters can be divided into two subclasses; DctP-type and TAXI type, which share the same overall architecture and requirement of the SBP for transport, but their SBPs share no similarity. The DctP-type transporters are very well studied and have been shown to transport a range of compounds including dicarboxylates, keto acids, sugar acids. However, the TAXI type transporters are relatively poorly understood, with the range of transportable compounds still to be discovered and selectivity requirements for binding unknown. To address these shortfalls in our understanding, we have structurally and biochemically characterized VC0430 from *Vibrio cholerae* revealing it to be a monomeric high affinity glutamate binding protein. VC0430 is stereoselective, binding the L-isomer preferentially, and can also bind L-glutamine and L-pyroglutamate, but with low affinity relative to L-glutamate. Structural characterization of ligand bound VC0430 reveals details of the binding site and biophysical characterization of binding site mutant reveal the substrate binding determinants, which differ substantially from the DctP-type TRAPs. Finally, we have analysed *in silico* the interaction between VC0430 and its cognate membrane component revealing an architecture hitherto unseen. To our knowledge, this is the first transporter in *V. cholerae* to be identified as specific to glutamate, which plays a key role in osmoadaptation of *V. cholerae*, making this transporter a potential therapeutic target.

## Introduction

The ability to take up extracellular solutes is essential for bacteria to thrive in an environmental niche or adapt to changing conditions. Prokaryotes have developed many transport systems that vary based on the substrates they transport, the energy source they use, the rate of transport required, and the affinity for their substrate. Many transporter families are secondary active transporters and utilize electrochemical gradients across the membrane (usually Na^+^ ions or protons). While most secondary active transporters are composed only of integral membrane protein components, some transporter families have recruited a substrate binding protein (SBP) that imparts directionality to transport and provides substrate affinities often in the nanomolar range, ideal for scavenging scarce nutrients^1^.

Tripartite ATP-independent periplasmic (TRAP) transporters are a large family of SBP-dependent secondary-active transporters that are found exclusively in prokaryotes to catalyze the uptake of a variety of substrates, primarily organic anions, across the cytoplasmic membrane^2,3^. A defining feature of TRAP transporters is the obligate utilization of an SBP, which is either found in the periplasm or attached to the outer leaflet of the cytoplasmic membrane, where it binds and delivers substrate to the membrane component. The membrane-embedded component of TRAP transporters consists of either two separate unequally sized integral membrane proteins, or a fusion of these two proteins into a single polypeptide with distinct domains^2^.

TRAP transporters can be divided in to two subtypes; the DctP-type, named after the first TRAP transporter characterized, which was specific to dicarboxylates^4^, and the TRAP-associated extracytoplasmic immunogenic (TAXI)-type, named due to the immunogenic properties of the first one identified in *Bacillus abortus*^5,6^. While the membrane components of each of these TRAP subtypes share substantial sequence identity suggesting a shared lineage and likely the same overall fold, the subtype-specific SBPs bear no relation to each other^6^.

The vast majority of our understanding of TRAP transporter structure and function comes from the characterization of DctP-type TRAP transporters^4,7–20^, most prominently, a sialic acid-specific member from *Haemophilus influenzae*, SiaPQM (HiSiaP is the SBP and HiSiaQM is the fused membrane component)^3,20–26^. Work from multiple labs on a variety of TRAP SBPs has revealed that, like SBPs from ATP-binding cassette (ABC) transporters, they utilize a venus flytrap type mechanism, in which the binding of ligand to the open *apo* state triggers closure of the two α/β globular domains around the substrate^27^. In addition, structural and phylogenetic analysis of DctP-type SBPs has revealed the presence of an extremely well conserved arginine residue in the binding site that makes a critical salt bridge with the ligand, which in almost all cases is an organic acid^26–28^. Functional reconstitution of SiaPQM from *H. influenzae, Photobacterium profundum* and *Vibrio cholerae* revealed that these transporters, and likely all DctP-type TRAPs use a Na^+^ gradient to power transport^19–22^.

While there are dozens of DctP-type SBP structures available revealing a broad range of substrates^1,2,16^, it was not until recently that experimentally derived structures of the membrane components were elucidated^19,20,24^. Structural analysis of HiSiaQM from *H. influenzae* and *P. profundum* has revealed important details regarding the interaction between the membrane component and the SBPs and the overall architecture of the membrane component. As previously predicted^29^, TRAP transporters share the same fold as members of the divalent anion Na^+^ symporter (DASS) family^30–33^, and both employ an elevator-like mechanism to transport substrate across the membrane^31,34,35^.

Compared to the DctP-type TRAPs, TAXI-TRAP transporters are relatively poorly studied despite being widespread in prokaryotes, and they differ in several ways. As with DctP-type TRAP transporters, TAXI-TRAPs are found in both bacteria and archaea, but TAXIs are the *only* type of TRAP transporter found in archaea, suggesting that TAXIs are the more ancient of the two subtypes^6^. In addition, while the DctP-type transporter membrane components can be composed of two separate polypeptides or a fusion of the two, TAXI-TRAP membrane components are only ever a single fused polypeptide. Finally, functional reconstitution of an α-ketoglutarate specific TAXI from *Proteus mirabilis* reveals that, instead of being driven by Na^+^ electrochemical gradient as for the characterized DctP-type TRAPs, this TAXI is powered by a H^+^ gradient, which is a significant mechanistic schism in the family^36^.

In contrast to DctP-type TRAPs, there are no published structures of TAXI membrane components, and there is only one published TAXI SBP structure, a glutamate or glutamine binding protein (TtGluBP) from *Thermus thermophilus*^37^. However, the identity of the bound substrate was ambiguous and there was no further investigation into binding determinants. There is also an unpublished structure of a TAXI SBP from *Erlichia Chaffeensis* (4DDD) in the PDB, which has a glycerol molecule and chloride ion bound in the vicinity of the binding site, but it is not clear whether these are substrates. A TAXI transporter from *Azoarcus* sp. CIB has been shown to transport orthophthalate^38^, and phenotypic analysis of a TAXI gene knockout in *Psychrobacter arcticus* suggest it is involved in the uptake of butyrate, glutamate, fumarate, and acetate^39^. This limited number of examples hints that TAXIs can also transport a range of substrates akin to the DctP-type TRAPs. To provide more insight into the substrate range, structural arrangement and key binding determinants of TAXI SBPs, we have obtained the structure of VC0430 from the human pathogen *V. cholerae*, determined the substrate specificity and affinity, and ascertained the effects of binding site mutation on ligand interactions. We have also investigated the interaction between the SBP and membrane components using modelling approaches revealing a hitherto unreported interfacial arrangement for any TRAP transporter.

## Methods

### Molecular Biology

Site-directed mutagenesis was performed using the Quikchange II kit (Agilent) or using KOD Hot-start DNA polymerase (Merck) followed by DpnI treatment. All plasmids and mutants were sequence verified prior to use.

### Expression of VC0430

To overexpress VC0430 for the periplasmic localization and ESI-MS analysis, the gene encoding VC0430 was expressed in BL21 (DE3) from a pET20b plasmid with the gene in-frame with a cleavable N-terminal PelB signal sequence and a C-terminal His_6_ tag. For all other analyses, the gene encoding VC0430 was overexpressed in BL21 (DE3) transformed with a pET-based plasmid with the gene in frame with an N-terminal FLAG/His_10_ tag (pETnHisVC0430) in place of the signal peptide (first 30 residues, MKEGKFMSLPKIIKMGAIAAAVIGSGVASA)^40^.

BL21(DE3) pETnHisVC0430 was grown in LB media supplemented with 50 µg/ml kanamycin and incubated at 37°C until an OD of 0.6-0.8 was reached, at which point expression was induced by addition of 1 mM IPTG. The induced cells were incubated overnight at 37°C, harvested by centrifugation at 4000 rcf for 20 min, resuspended in Purification Buffer (PB, 50 mM Tris, pH 8, 200 mM NaCl, 5% v/v glycerol), and lysed by 3 passes through a cell disruptor (Avestin). The lysate was clarified by centrifugation at 20000 xg for 20 min at 4°C. BL21(DE3) pET20bVC0430 was treated similarly, except cells were grown at 25°C, cells were induced at and OD of 0.4 and cells were lysed using a French pressure cell.

### Purification of VC0430

The periplasmically located VC0430 that was used for ESI-MS analysis was purified by applying the clarified lysate to N-NTA resin (Qiagen), washing the resin with Wash Buffer (WB, PB containing 20 mM imidazole), and then eluting bound VC0430 with PB containing 300 mM imidazole. Eluted protein was further purified using size exclusion chromatography SEC) prior to analysis. N-terminally FLAG/His-tagged VC0430, which was used for all other analysis, was denatured during the purification process to remove any pre-bound ligand. Clarified lysate was incubated with Ni-NTA resin, which was then washed with 30 column volumes (CV) of WB + 2 M guanidinium chloride (GdmCl) to denature the protein. Protein was refolded by subsequent washes with 4 CV WB + 1.5 M GdmCl, 4 CV WB + 1 M GdmCl, 4 CV WB + 0.5 M GdmCl and finally 8 CV WB. Refolded protein was eluted with PB + 300 mM imidazole. The FLAG/His tag was cleaved for binding analysis and crystallography by incubation with TEV protease for 4 hours at room temperature before reapplying the mixture to Ni-NTA resin and collecting the tag-free VC0430 in the flowthrough. For the protein used for crystallography, VC0430 was further purified using size exclusion chromatography with an Superdex 200 Increase 10/300 GL column (GE Healthcare).

### Oligomeric state analysis using SEC standards

For molecular weight measurements, Ni-NTA purified VC0430 was concentrated and applied to an Superdex 200 Increase column at a flowrate of 0.5 ml/min with SEC buffer (50 mM Tris, pH 7.5). The calibration curve was generated by analysing SEC standards (Thermo Scientific) under than same conditions as VC0430. The partition coefficient (K_av_) for the standards and VC0430 using the equation: K_av_ – (V_e_-V_o_)/(V_t_-V_o_), where V_e_ is the elution volume, V_o_ is the column void volume, and V_t_ is the total column volume.

### Differential scanning fluorimetry (DSF)

To perform DSF, 5 µM protein was mixed with substrate (0, 1, or 15 mM) and 2.5x SYPRO Orange dye (ThermoFisher), and made up to a total volume of 50 µl with DSF buffer (50 mM Tris pH 8, 20 mM NaCl). Using a QuantStudio 3 RT-PCR thermocycler (Invitrogen), the DSF samples were incubated at 5°C for 1 min, then the temperature increased to 95°C in 1°C increments, holding at each temperature for 10 s. The reporter dye setting used was set to SYBR. Melt curve data were exported to Microsoft Excel and GraphPad Prism for analysis and presentation.

### Tryptophan fluorescence spectroscopy

All tryptophan fluorescence assays were performed using a Cary Eclipse fluorimeter (Agilent). Fluorescence emission spectra were collected using 0.5 µM VC0430 in 50 mM Tris, pH 7.4, at 20°C, exciting at 295 nm wavelength with an emission range of 300-400 nm, slit widths set to 10 nm, and medium PMT voltage. Data were smoothed using a Savitzky-Golay smoothing factor of 15 for the initial emission scans. Ligand titrations were performed using time-based acquisition with λex of 295 nm, λem of 330 nm, excitation and emission slit widths of 5 and 10 nm, respectively. Following baseline fluorescence collection for 30 s, the fluorescence change for each addition of ligand was measured for 20s, which was averaged to calculate the Δfluorescence for each addition. Binding curves were analysed using GraphPad prism and Kd values were obtained by fitting the curves to a one site total binding model.

### Mass spectrometry

For mass spectrometry, SEC-purified VC0430 was dialysed against 50 mM sodium phosphate, pH 8, dialysed against water, and then concentrated prior to analysis. Electrospray mass spectrometry was performed using the API Qstar mass spectrometer using an ionspray source. To determine the mass for the unliganded protein, 1 µM VC0430 was made up in acetonitrile and 0.1% formic acid. To determine the mass of the protein with bound ligand, 10 µM VC0430 was made up in 25 mM ammonium acetate, pH 4.5 and 3% (v/v) methanol. Raw m/z data were deconvoluted using Bayesian Protein Reconstruction routine.

### Protein crystallography

VC0430 (15 mg/ml final concentration) was incubated with 0.5 mM L-glutamate before being mixed in a 1:1 ratio with 2.4 M sodium malonate at pH 7.0 and (JCSG+ Condition F9 with no changes made to the solution), over a well of the same buffer in a hanging drop vapor diffusion plate. For x-ray diffraction, the crystals were then picked using a 0.3-0.4 µm nylon loop. The crystals were immediately flash frozen in liquid nitrogen. X-ray diffraction experiments were conducted at Diamond Light Source using the macromolecular beam i04 with a wavelength of 0.9537 Angstroms. Data was collected over a 360° rotation of the omega axis, collecting images at 0.1 degrees. The full statistics for the crystals can be found in Supplementary Table 2. In brief, the crystals diffracted to approximately 1.7 Angstroms with a CC1/2 of 1.0 over the data range. The crystals had a space group of R32 and the data collected were used to solve the structure using the Diamond Light Source automated pipeline. The data were indexed, integrated and scaled by the by xia2 3dii pipeline^41^. The processed data was then used for molecular replacement model obtain through AlphaFold2^42^) and autobuilding using Dimple^43^. The initial structure consisted of 300 built-in residues; additional building, additional residue and water placement was performed using Coot; and Phenix was used for refinement and automated water placement^44,45^.

### Bioinformatics

A total of 59 TAXI amino acid sequences from a wide range of species were obtained from Uniprot and aligned using MAFFT from EMBL-EBI^46^. The alignment was visualized in Geneious Prime 2022.2.1 (https://www.geneious.com) and consensus residues were highlighted. To construct the maximum-likelihood phylogenetic tree, alignment gaps at the N- and C-terminus were trimmed. A phylogenetic tree was created using IQ-TREE with 1,000 replicates of ultra-fast bootstrapping approximation^47^. The unrooted consensus tree was then visualized using the Interactive Tree of Life (iTol) tool^48^.

## Results

### Bioinformatic analysis of VC0430

To characterize VC0430, we first sought to identify its ligand. As a starting point, we analyzed the genome context of *VC0430*, to see if characterized co-regulated genes could provide clues to the identity of the ligand; a strategy used to much success previously^9,15^. As expected, the gene encoding VC0430 is adjacent to the gene encoding its cognate membrane component, Vc0429 (Fig. 1A). The TRAP transporter genes are downstream of 2 other genes, *argR* and *mdh* which encode an arginine repressor and malate dehydrogenase, respectively (Fig. 1A)^49^. Arginine and malate both contain carboxyl groups, which is a common feature of known TRAP ligands, raising the possibility that they are substrates of the transporter, but the lack of support for coregulation between these genes and the TRAP genes diminishes the strength of this prediction. The gene downstream of the TRAP genes encodes universal stress protein (UspA); the association between TRAP transporters and UspA has been noted previously^2,50^, and while they appear to influence TRAP transporter function, their physiological role and mechanism are not known^50^.

**Figure 1.**
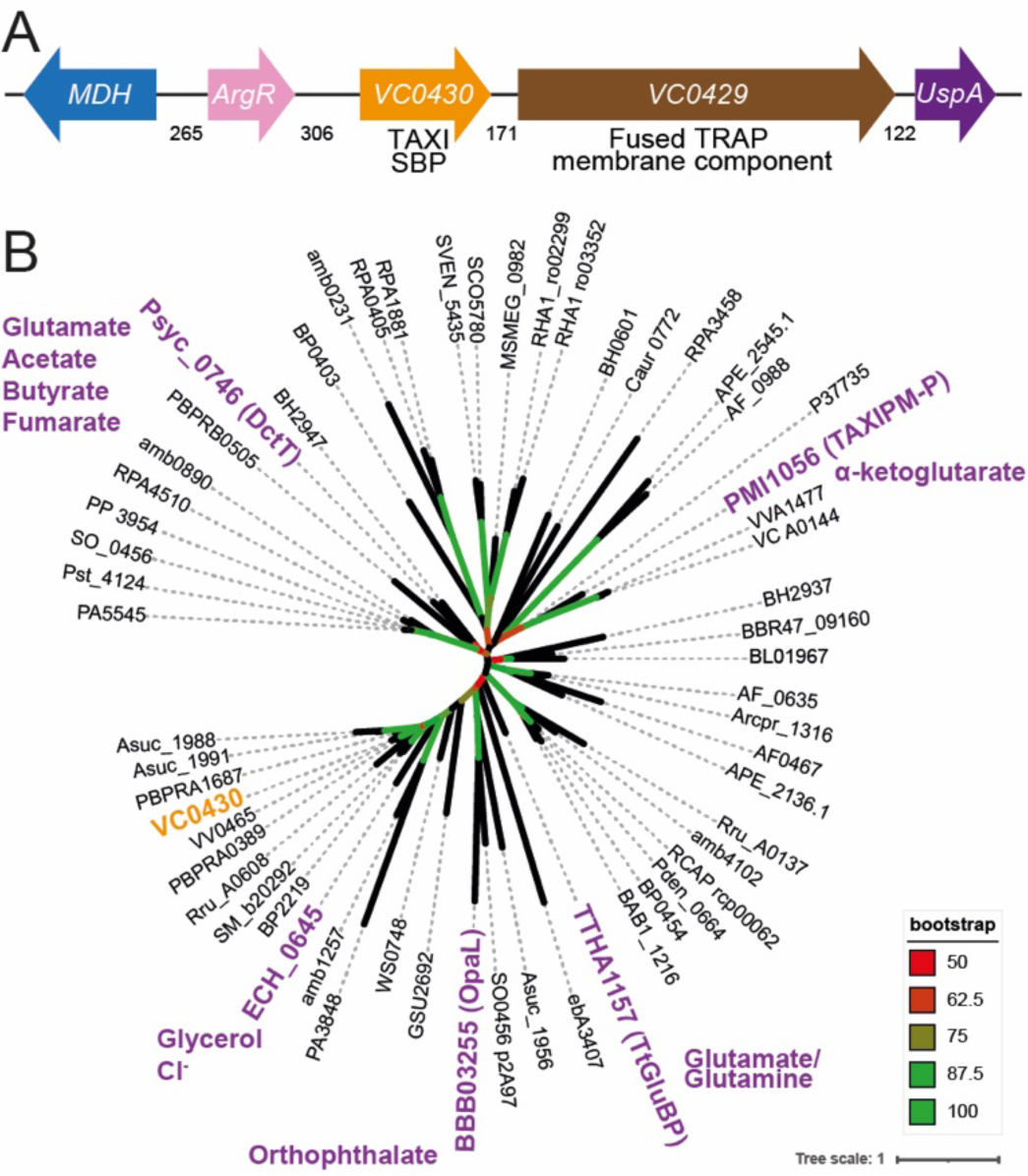
Genome context and phylogenetic analysis of VC0430. **A)** Genome context of *VC0430*. Numbers indicate the number of intergenic basepairs. **B)** Phylogenetic tree of 59 homologues of VC0430 (locus tag for each protein is displayed). The TAXI SBPs that have been characterized to any degree have been highlighted in purple and the known or predicted substrate(s) is indicated. VC0430 in orange is the subject of this study. The accompanying sequence alignment is presented in Supplementary figure 1.

An alternative approach to predict the substrate of orphan transporters is to see if any closely related SBPs are characterized because they may bind the same or related ligands. To perform this analysis, we collected and aligned the amino acid sequences of 59 TAXI SBPs from a variety of organisms, making sure to include any characterized TAXI SBPs (Fig. 1B). The phylogenetic tree produced by this alignment revealed several clades. However, when the positions of the TAXIs with known or predicted substrates were mapped onto the tree, none were sufficiently closely related to VC0430 to permit any predictions (Fig. 1B). In the absence of strong evidence for the identity of VC0430’s ligand from bioinformatic approaches, we sought to identify the ligand experimentally. As SBPs generally have a high affinity for their cognate ligand, SBPs can retain the bound ligand during the purification process if that ligand is present in the expression strain. Therefore, we decided to identify VC0430’s ligand by expressing and purifying the protein and screening for the cognate ligand using mass spectrometry, an approach used previously to de-orphanise SBPs^9,15,16^.

### VC0430 is monomeric and interacts with both glutamate and glutamine

To identify any ligands bound to VC0430 using this mass spectrometry approach, we overproduced VC0430 in *E. coli* BL21 (DE3) with an N-terminal cleavable signal peptide and a C-terminal His_6_-tag, which would translocate the protein to the periplasm, and facilitate purification, respectively. We purified VC0430 in one step with immobilized metal affinity chromatography (IMAC) and subjected to it to mass spectrometry analysis under denaturing conditions, revealing a mass of 33573.9 Da, which correlates well with the predicted mass of the mature protein with the C-terminal affinity tag (33578 Da, Fig. 2A). While the vast majority of structurally characterized TRAP SBPs are monomeric, there are some notable exceptions that form stable dimers^51^. However, the close correlation of the mass of denatured VC0430 with the predicted mass indicates that VC0430 is monomeric. Under more native conditions, we observed 2 major peaks in the mass spectrum, the first with a mass of 33573.5 which is essentially identical to the mass for the denatured protein suggesting it corresponds to the *apo* protein (Fig. 2B). The second major peak had a mass of 33719.7, which we reasoned corresponded to ligand bound VC0430 (Fig. 2B). Calculating the difference between the 2 major MS peaks under native conditions revealed a mass difference of 146.2 (±2 Da), providing us with an approximate mass of the bound ligand; an important clue in its identification. Two smaller peaks were present that were spaced 98 Da from each of the main peaks suggesting that they are phosphate adducts (Fig. 2B).

**Figure 2.**
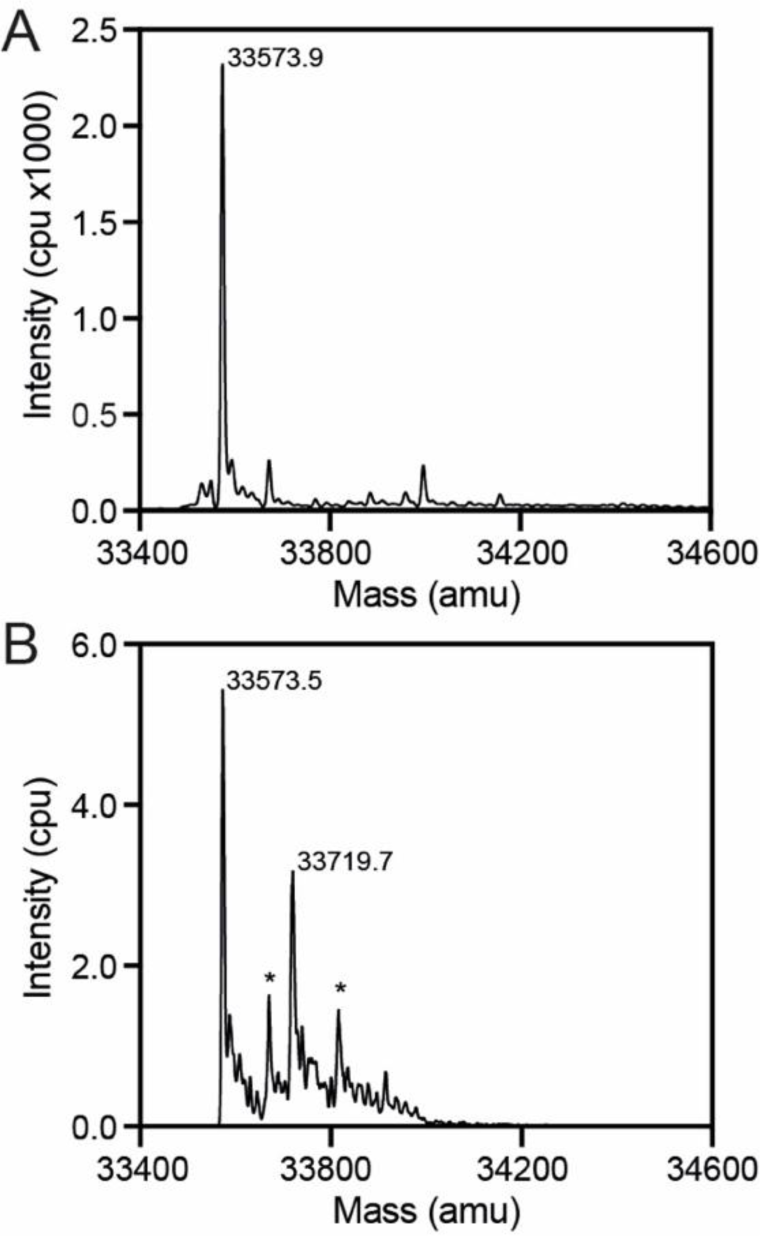
Identification of ligand mass using mass spectrometry. **A)** Denaturing MS analysis of VC0430 reveals a single peak corresponding to a mass of 33573.9 Da. **B)** Native MS analysis of VC0430 reveals major peaks corresponding masses of 33573.9 and 33719.7, which represent the *apo* and ligand bound VC0430, respectively. Peaks labelled with an * correspond with phosphate adducts of the major peaks.

Armed with the approximate mass of the ligand (144-148 Da), we narrowed the list of potential compounds further by restricting our search to metabolites present in *E. coli* using the EcoCyc^52^. This search resulted in a list of 51 different compounds (Supplementary table 1).

To increase our yield of VC0430 for structural and biochemical studies, we changed expression construct and replaced the N-terminal signal peptide with a TEV-cleavable decahistidine tag, thus leading to cytoplasmic expression. VC0430 was overexpressed using this new construct and purified using IMAC and size exclusion chromatography (SEC), which revealed a single band in SDS-PAGE and a sharp, symmetrical peak in the SEC, indicative of well folded and stable protein (Fig. 3A). We compared the elution volume of VC0430 from the SEC column to standards with known molecular weights (Fig. 3A). Analysis of the calibration curve revealed that VC0430 has a mass of 28.2 kDa (Fig. 3B), which is slightly smaller than the molecular weight predicted from the amino acid sequence (35.9 kDa), but it demonstrates that VC0430 exists as a monomer in solution, strongly supporting our mass spectrometry data.

**Figure 3.**
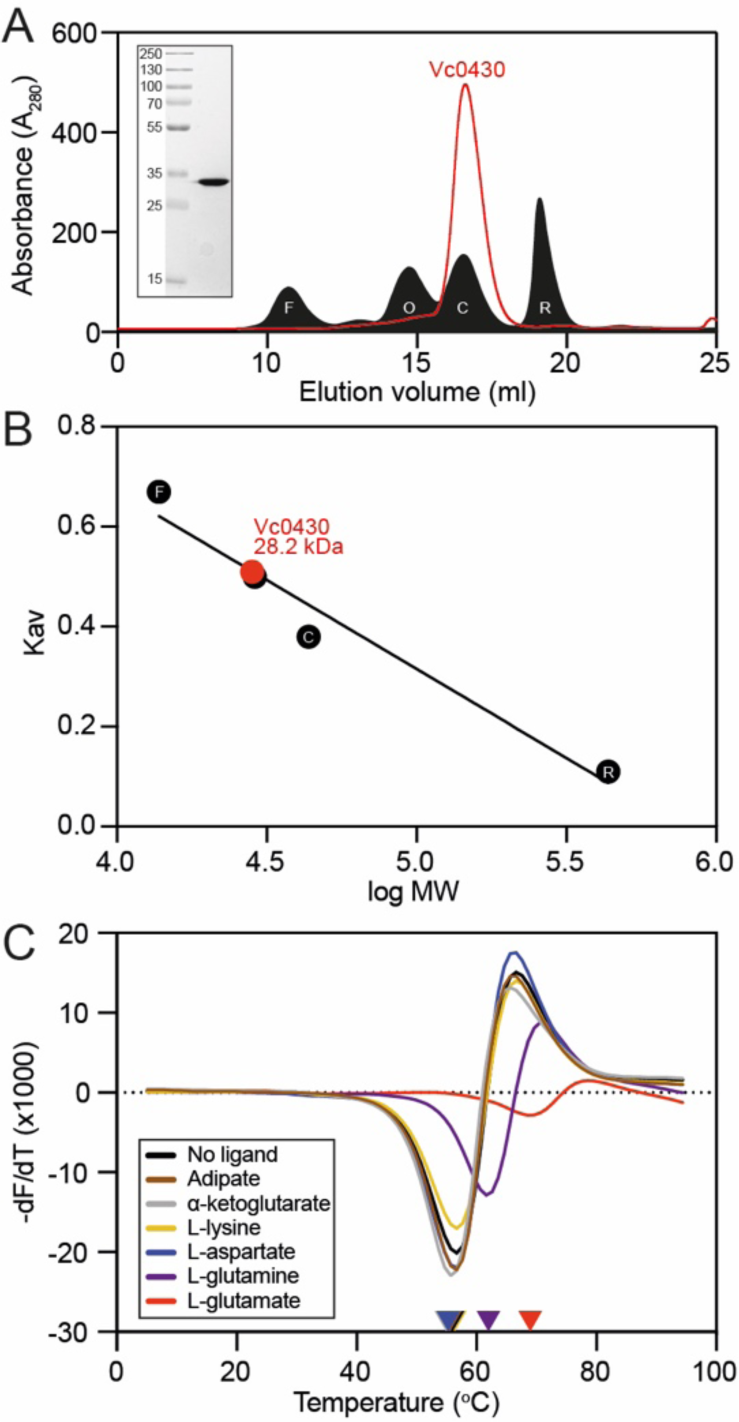
Purification, analytical size exclusion chromatography and ligand binding of VC0430. **A)** SEC elution profile of VC0430 (red line) compared to the profiles of 4 standards (filled black trace); ferritin (F, 10.67 ml elution volume), ovalbumin (O, 14.71 ml), carbonic anhydrase (C, 16.52 ml), and ribonuclease A (R, 19.08 ml). **B)** Molecular weight calibration curve showing Kav as a function of the log MW of the 4 standards from **A** (black circles, labels are the same as in **A**). Comparison of the VC0430’s Kav reveals a MW of 28.2 kDa. **C)** Derivatives of the unfolding curves (dF/dT) for VC0430 in the absence of ligand and the presence of 1 mM adipate, a-ketoglutarate, L-aspartate, L-glutamate, L-glutamine and L-lysine. Coloured arrow on the X-axis indicate the apparent protein Tm under those conditions.

To screen our list of compounds to see which could bind to VC0430, we used differential scanning fluorimetry (DSF). In this assay, protein is thermally denatured in the presence of SYPRO Orange dye, the fluorescence of which increases as it binds to the hydrophobic core of the protein that is revealed during denaturation. DSF provides a read-out of the melting temperature (Tm) of the protein, which often increases upon ligand binding, thus providing a convenient way of detecting interactions.

Using DSF, we first screened the most conveniently available compounds from our EcoCyc-derived list of potential ligands; adipate, α-ketoglutarate, L-aspartate, L-glutamate, L-glutamine and L-lysine. Analysis of the first derivative of the fluorescence as a function of the temperature (dF/dT) for VC0430 alone revealed a Tm of 57.6°C (Fig. 3C). In the presence of 1 mM L-aspartate, L-lysine, α -ketoglutarate or adipate, we observed no change in VC0430’s Tm suggesting that these compounds do not bind (Fig. 3C). However, upon addition of 1 mM L-glutamate and L-glutamine, we observed a substantial rightward-shift of the melt peak, with a Tm increase of 11.3°C and 5.6°C, respectively, indicating that both L-glutamate and L-glutamine bind to VC0430 (Fig. 3C).

### Determining the binding affinity of VC0430

To provide support for L-glutamate and L-glutamine being ligands of VC0430, we determined VC0430’s binding affinity for each compound using intrinsic tryptophan fluorescence. When excited at 295 nm, VC0430 exhibited a characteristic tryptophan emission spectrum between 300-400 nm with an emission maximum of ∼330 nm (Fig. 4A and B, black line). Upon addition of 5 µM L-glutamate we observed a substantial 7.4% enhancement of the fluorescence at 330 nm, and a smaller 4.9% enhancement upon addition of 100 µM L-glutamine, indicative of binding (Fig. 4A and B, red line). To determine the binding affinities for L-glutamate and L-glutamine, we monitored the dose-dependent enhancement of fluorescence in the presence of increasing concentrations of each ligand (Fig. 4A and B, inset). Fitting the curves to one site binding model revealed dissociation constants (Kd) of 0.10 ± 0.05 µM and 18.53 ± 6.78 for L-glutamate and L-glutamine, respectively. These Kd values are consistent with the affinities measured for other DctP-type TRAPS SBPs and the other TAXI SBP that has been characterized^2,36^. As these data suggest that L-glutamate is the preferred ligand for VC0430, we propose that VC0430 should be designated VcGluP (with the cognate membrane component Vc0429 designated VcGluQM), to remain consistent with the naming of DctP-type TRAP transporters.

**Figure 4.**
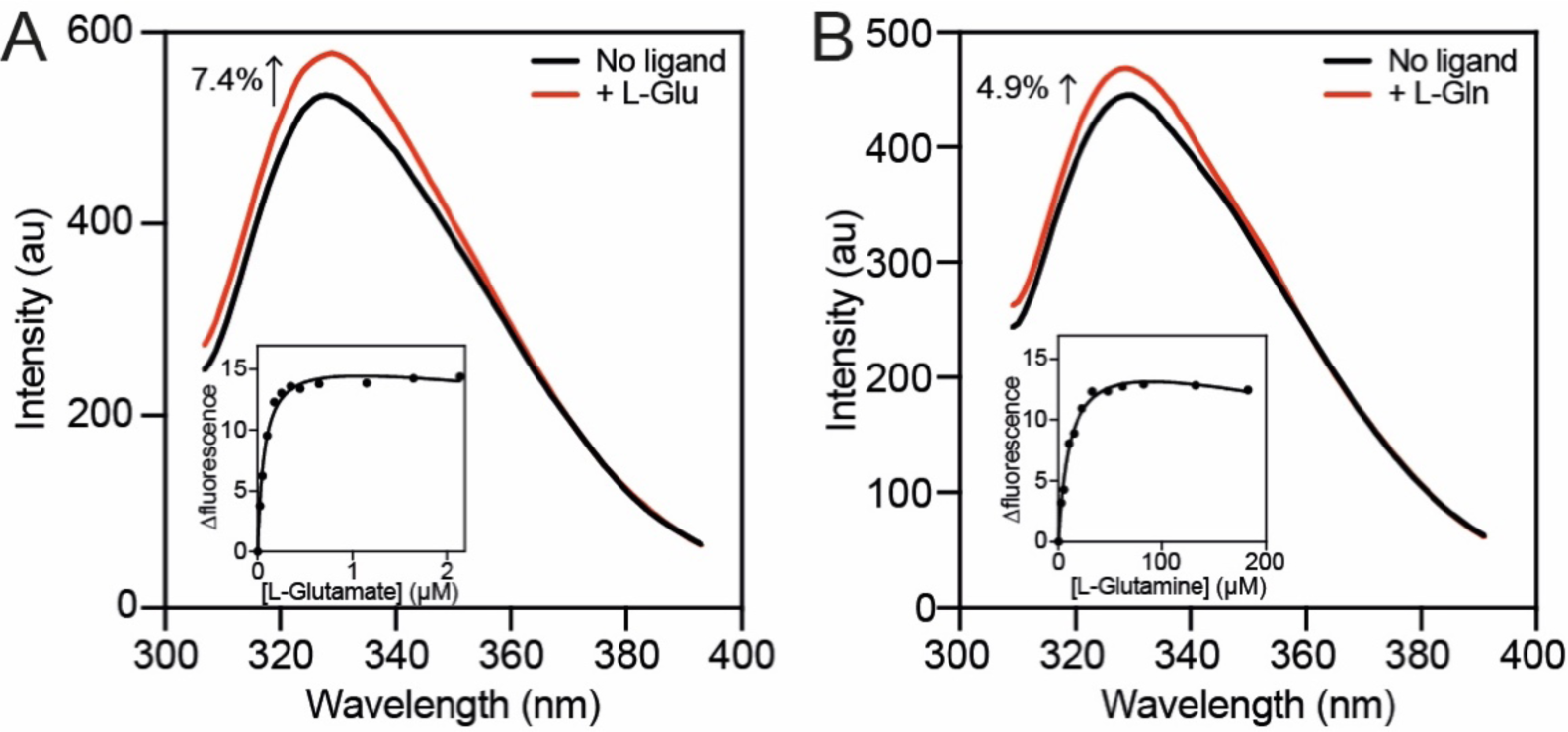
Ligand binding affinity determination using intrinsic tryptophan fluorescence. Fluorescence emission scans of VcGluP in absence of ligand (black lines) and in the presence of **A)** 5 µM L-glutamate and **B)** 100 µM L-glutamine (red lines). *Inset:* representative L-glutamate (A) and L-glutamine (B) binding curves for VcGluP showing hyperbolic dose response. Single data sets are shown, but assays were performed at least 3 times with equivalent outcomes.

### VcGluP stereoselectively binds its ligands

Having identified that VcGluP binds both L-glutamate and L-glutamine, we next assessed VcGluP’s stereoselectivity for these ligands by evaluating its ability to bind D-glutamate and D-glutamine. DSF analysis revealed that, while L-glutamate induces a sizeable 11.6°C increase in thermostability of VcGluP, 1 mM D-glutamate induces lower but above background stabilization of 4.0°C (Fig. 5A and C). D-glutamine has minimal effects on VcGluP stabilization, only inducing ∼1.7°C stabilization upon addition of 15 mM (Fig. 5B and C). To provide further insight into the stereoselectivity, we determined the binding affinity for the D-enantiomers using tryptophan fluorescence revealing a Kd of 24.5 ± 10.6 µM for D-glutamate (Fig 5D). However, a Kd for D-glutamine could not be determined due to lack of fluorescence enhancement. These data demonstrate that L-glutamate is VcGluP’s preferred ligand, but it is still able to bind the D enantiomer with an affinity in the range seen for other SBPs.

**Figure 5.**
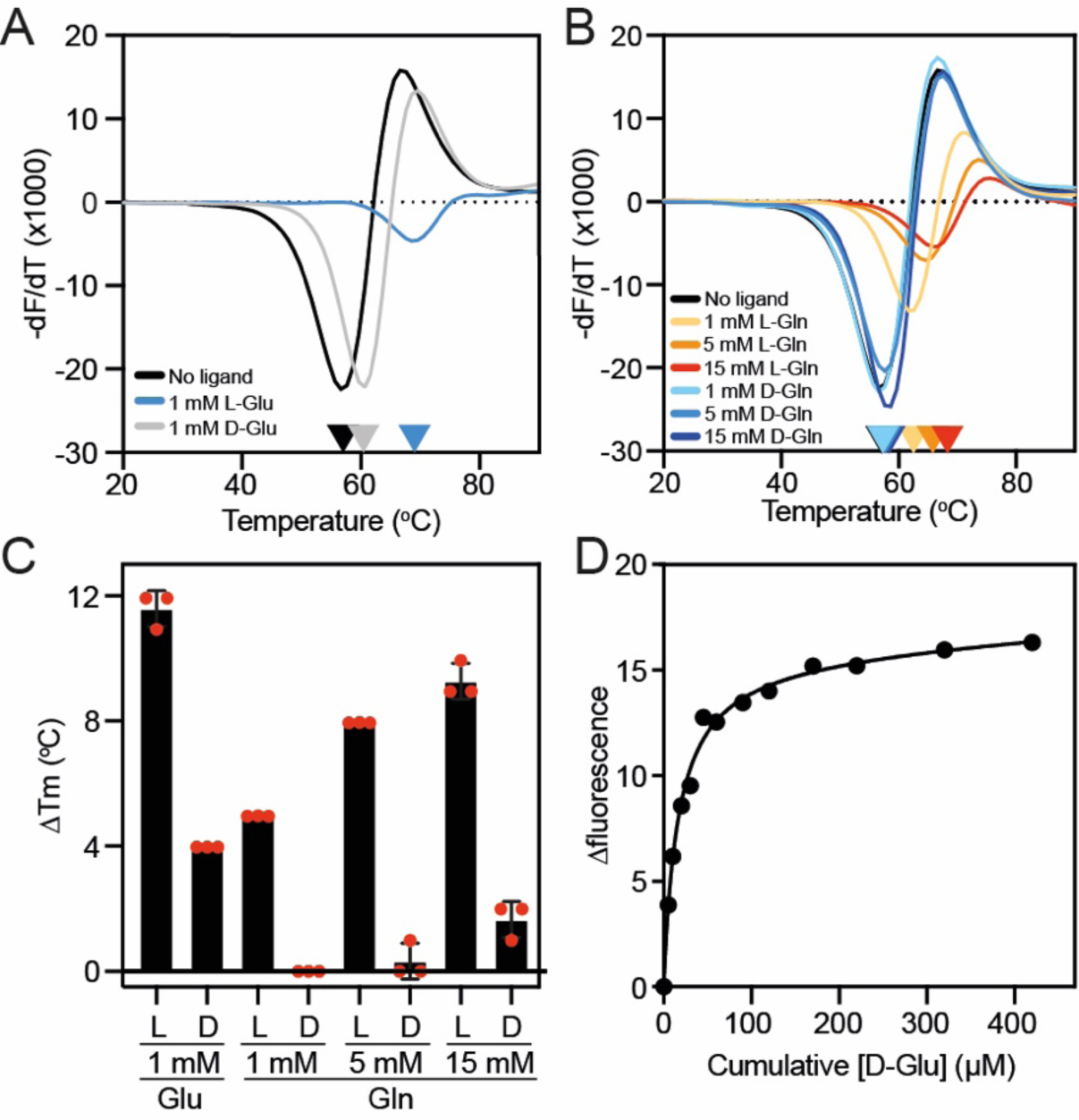
Assessing the stereoselectivity of VcGluP. Derivatives of the unfolding curves (dF/dT) for VC0430 in the absence of ligand and the presence of **A)** 1 mM L- and D-glutamate, and **B)** 1, 5 and 15 mM L- and D-glutamine. **C)** Thermostabilisation of VcGluP in the presence of L- and D-glutamate and glutamine. Error bars represent standard deviation and individual data points are shown as red circles. **D)** Representative binding curve for D-glutamate binding to VcGluP.

### L-pyroglutamate is a low affinity ligand of VcGluP

Due to the high affinity for L-glutamate exhibited by VcGluP and evidence of 4 other transporter-related SBPs binding the similar molecule L-pyroglutamate^13,16^, we tested whether VcGluP could also bind L-pyroglutamate. DSF screening revealed that addition of L-pyroglutamate enhanced VcGluP’s stability in a dose-dependent manner (Fig. 6A), suggesting an interaction. In support of this, we observed 8% enhancement in VcGluP’s intrinsic tryptophan fluorescence upon addition of 1 mM L-pyroglutamate, which did not increase upon further additions, suggesting saturable binding (Fig. 6B). We investigated the affinity of L-pyroglutamate binding using tryptophan fluorescence revealing a Kd of 73.54 ± 27.93 µM (Fig. 6C). Therefore, while the affinity for L-pyroglutamate is >700x lower than the affinity for L-glutamate, VcGluP can clearly bind L-pyroglutamate, which is surprising considering the pronounced chemical differences between the two ligands (Fig. 6D).

**Figure 6.**
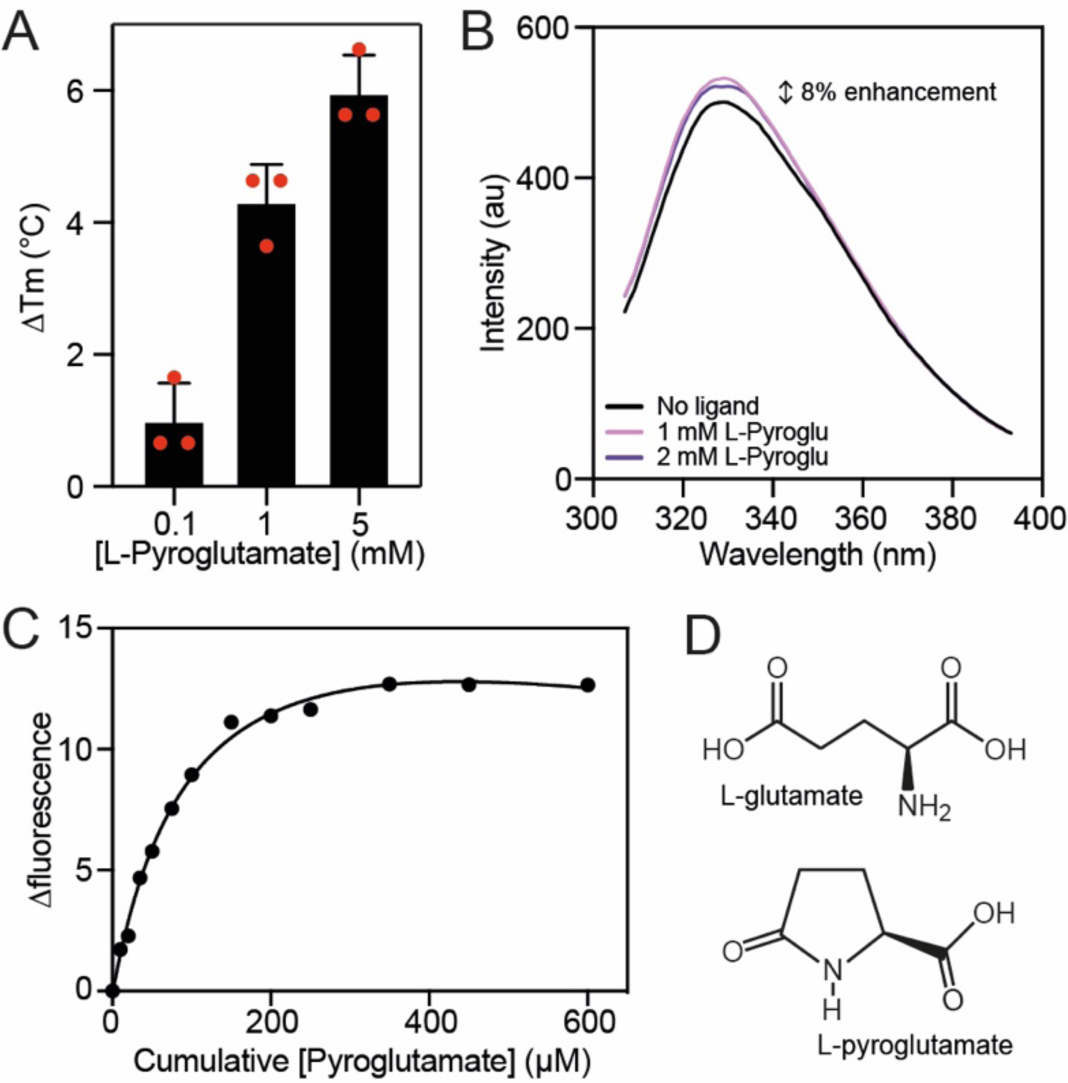
VcGluP can also bind L-pyroglutamate. **A)** Thermostabilisation of VcGluP in the presence of 0.1, 1 and 5 mM L-pyroglutamate. Error bars represent standard deviation and individual data points are shown as red circles. **B)** Fluorescence emission scans of VcGluP in the presence and absence of L-pyroglutamate. **C)** Representative binding curve for L-pyroglutamate binding to VcGluP. **D)** Chemical structures of L-glutamate and L-pyroglutamate.

### Structural characterization of L-glu bound VcGluP reveals binding site residues

To investigate the selectivity determinants of VcGluP, we solved the structure of VcGluP in the presence of L-glutamate to 1.7 Å, using molecular replacement (Fig. 7a). The refined model contains residues 29-328, which equates to all of the 299 residues of mature VcGluP. VcGluP is composed of 2 α/β domains connected by a hinge composed of antiparallel β-sheets (residues 116-131 and 255-267). Domain 1 is composed of amino acids 29-115 and 268-328, and domain 2 consists of residues 132-254. Each domain contains 3 central parallel β-sheets bracketed by α-helices. Comparison with other SBP structures reveals that VcGluP belongs to Cluster F^53^. In addition, each domain contains a disulfide bond; Cys53-Cys67 in domain 1 which connects a1 with B2, and Cys191-Cys216 in domain 2 that connects a8 and a9, although density for this disulfide is missing likely though radiation damage (Fig. 7C). We also identified 3 Na^+^ ions bound to the surface of the protein (Fig. 7, purple spheres), although due to their distance from the binding site, we predict that these are functionally irrelevant interactions and merely a consequence of the 2.4 M sodium malonate used during crystallisation.

**Figure 7.**
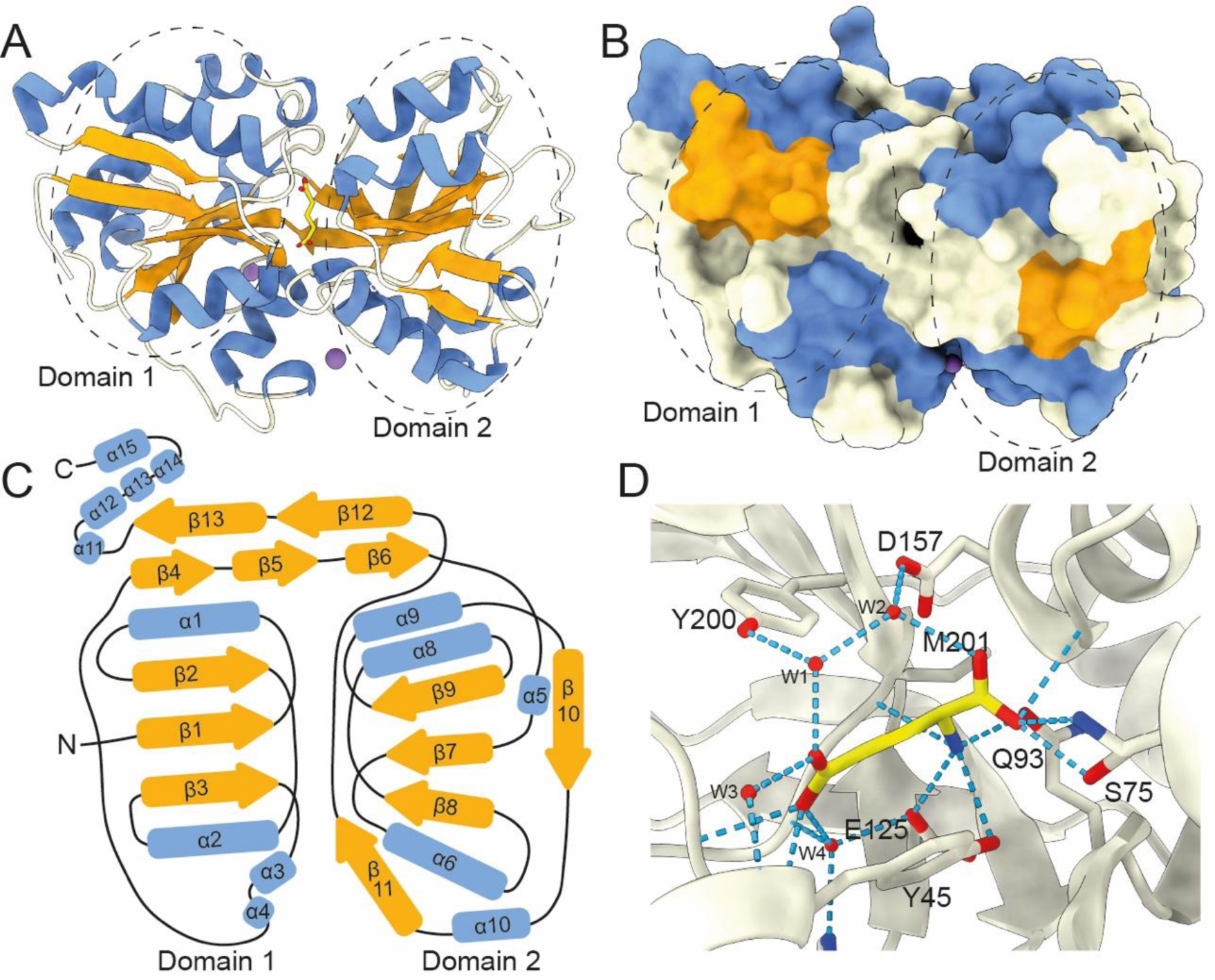
Structural characterisation of L-glutamate bound VcGluP. **A)** Cartoon representation of VcGluP coloured according to secondary structure, α-helices in blue, β-sheets in orange, loops in white. The bound L-glutamate is shown in yellow in the central cavity and the Na^+^ ions are purple spheres. Domains 1 and 2 are demarcated by dashed lines. **B)** Surface representaiton of VcGluP coloured and in same pose as in A). The lack of solvent accessibility to the binding site indicates VcGluP is in the closed state. **C)** Topology map of VcGluP, coloured as in A). **D)** Ligand binding site of VcGluP showing the interactions with the bounds L-glutamate.

During refinement, we identified non-protein density in the cleft between the two domains, which we assign to L-glutamate, which was present at 0.5 mM during crystallization (Fig. 7). As expected, in the presence of ligand, VcGluP adopts a closed conformation wherein the two domains completely envelope the ligand binding site (Fig. 7B).

Analysis of the ligand binding site reveals that L-glutamate is coordinated by VcGluP via a hydrogen bonding network composed of direct mainchain and sidechain interactions, and indirectly through coordinated water molecules (Fig. 7D). The α-carboxyl group of the bound L-glutamate makes sidechain and mainchain interactions with S75, mainchain interactions with G156 and hydrogen bonds to a water molecule, which is itself coordinated by the D157 sidechain. The amino group is coordinated by hydrogen bonding interactions with the side chains of Y45, Q93 and E125, and the mainchain of M201. The side chain carboxyl of the ligand makes mainchain interactions with V44 and Y45, and hydrogen bonds to several water molecules that are coordinated by S40, Y200, Y46, H123, G203 and E125. The positioning of Y45 and Y200 either side of the binding site likely also provides additional hydrophobic interactions with the ligand.

### Biochemical characterisation of binding site residue contributions

To investigate the binding determinants of VcGluP, we mutated the binding site residues Y45, S75, Q93, E125, Y200, M201 to alanine, expressed and purified them, and assessed the impact of these mutations on ligand binding using DSF and tryptophan fluorescence. Each VcGluP variant was overproduced and purified, and their stability was initially assessed using DSF. While there was some variation in Tm under *apo* conditions for the different variants, the presence of a characteristic melt curve suggested they were all folded and stable (Supplementary figure 2A). We next used tryptophan fluorescence emission scans to screen for ligand binding by the mutants (Supplementary figure 2B). Addition of L-glutamate induced enhancement of fluorescence for all the mutants indicating that all of them can bind ligand. However, while maximum fluorescence enhancement could be achieved with only 50 µM L-glutamate or less for Y45A, S75A, E125A, Y200A and M201A mutants, a higher concentration was required to saturate Q93A, indicating diminished affinity for this mutant (Supplementary figure 2B).

Using the dose-responsive fluorescence change for each mutant, we determined their Kds for the primary ligand L-glutamate using tryptophan fluorescence. Our titrations revealed that all 6 mutants exhibited a reduced affinity for L-glutamate compared to wildtype (Table 1). The most deleterious mutations were Y45A and Q93A, which reduced affinity 158 and 749-fold, respectively, followed by S75A, which had 10-fold lower affinity. E125A, Y200A and M201A had relatively modest effects on affinity, reducing it by ∼3-fold or less (Table 1).

**Table 1.** Equilibrium binding affinities for VcGluP wildtype and binding site mutants. FC is the fold change in Kd compared to wildtype, n indicates the number of replicates performed, and error indicated is the standard deviation. Values are colour coded to indicate the extent of Kd change for a mutant compared to wildtype (red being most deleterious, blue being a mild effect, and orange being middling).

| Variant | Glu $K_d$ (μM) | FC | n | Gln $K_d$ (μM) | FC | n | Pyroglu $K_d$ (mM) | FC | n |
| --- | --- | --- | --- | --- | --- | --- | --- | --- | --- |
| WT | 0.10±0.05 | - | 11 | 18.53±6.79 | - | 6 | 0.07±0.03 | - | 10 |
| Y45A | 15.82±1.58 | 158 | 5 | 8665.33±2603.74 | 468 | 6 | 2.87±0.09 | 41 | 5 |
| S75A | 0.99±0.14 | 9.9 | 5 | 96.07±38.94 | 5 | 7 | 0.65±0.06 | 9.2 | 5 |
| Q93A | 74.93±17.78 | 749 | 10 | ND | ND | 5 | 30.60±10.20 | 428 | 5 |
| E125A | 0.27±0.11 | 2.7 | 6 | 36.75±13.91 | 2 | 9 | 0.18±0.013 | 2.6 | 6 |
| Y200A | 0.10±0.47 | 0 | 8 | 23.57±8.78 | ND | 8 | 0.12±0.02 | 1.7 | 3 |
| M201A | 0.33±0.10 | 3.3 | 5 | 48.64±15.69 | 2.6 | 9 | 0.36±0.01 | 5.1 | 4 |

We also used tryptophan fluorescence to determine the mutants’ affinities for L-glutamine and L-pyroglutamate and observed the same pattern with Y45A and Q93A reducing the affinity the most, and E125A, Y200A and M201A having relatively little to no effect. Interestingly, the fold change in affinity is essentially the same for each mutant’s interaction with the 3 ligands, aside from Y45A where we observed 468-fold change for L-glutamine, but only 158-fold change for L-glutamate and 41-fold change for L-pyroglutamate (Table 1).

### Modelling the VcGluP-VcGluQM interaction reveals novel claw domain

There are currently no experimentally derived structures of TAXI membrane components, so to investigate the interaction between VcGluP with its cognate membrane component VcGluQM, we obtained the model of VcGluQM using Alphfold2^42^. The model produced by Alphafold2 shared many core features identified in the cryo-EM structure of HiSiaQM^24^, which is a membrane component from a DctP-type TRAP transporter. Both proteins consist of a transport or ‘elevator’ domain and the clearly separable scaffold or ‘stator’ domain (Fig. 8A and B). Most of the structural differences in these core domains appear to be localized to the stator, with the VcGluQM stator predicted to be somewhat larger than the one in HiSiaQM (Fig. 8A and B). However, there is substantial structural conservation in the transport domains, which superimpose very well with an RMSD of ∼4 Å over 230 residues (Fig. 8C). These structural similarities may not be surprising because both proteins belong to the TRAP transporter family. However, they share a remarkably low amino acid sequence identity of 7.9% (calculated using AlignMe^54^). In addition to the low level of identity, VcGluQM is 221 amino acids longer than HiSiaQM, the majority of which comes from a 185 amino acid C-terminal extension in VcGluQM. This extension (which we will refer to as the ‘claw’ domain) folds into a globular domain belonging to the uncharacterized DUF3394 protein family, is located on the periplasmic side of the membrane, and is mainly composed of β-sheets (Fig. 8). The positioning of the claw domain suggests that it likely comes into direct contact with VcGluP. To investigate this interaction, we modelled the holocomplex VcGluPQM using Colabfold^55^, which revealed that, as with the models of HiSiaPQM^24^, the SBP forms multiple interactions with the membrane component (Fig. 9). However, the model of VcGluPQM suggests that the C-terminal claw domain also plays a role in stabilizing the docking of the SBP with the membrane component (Fig. 9).

**Figure 8.**
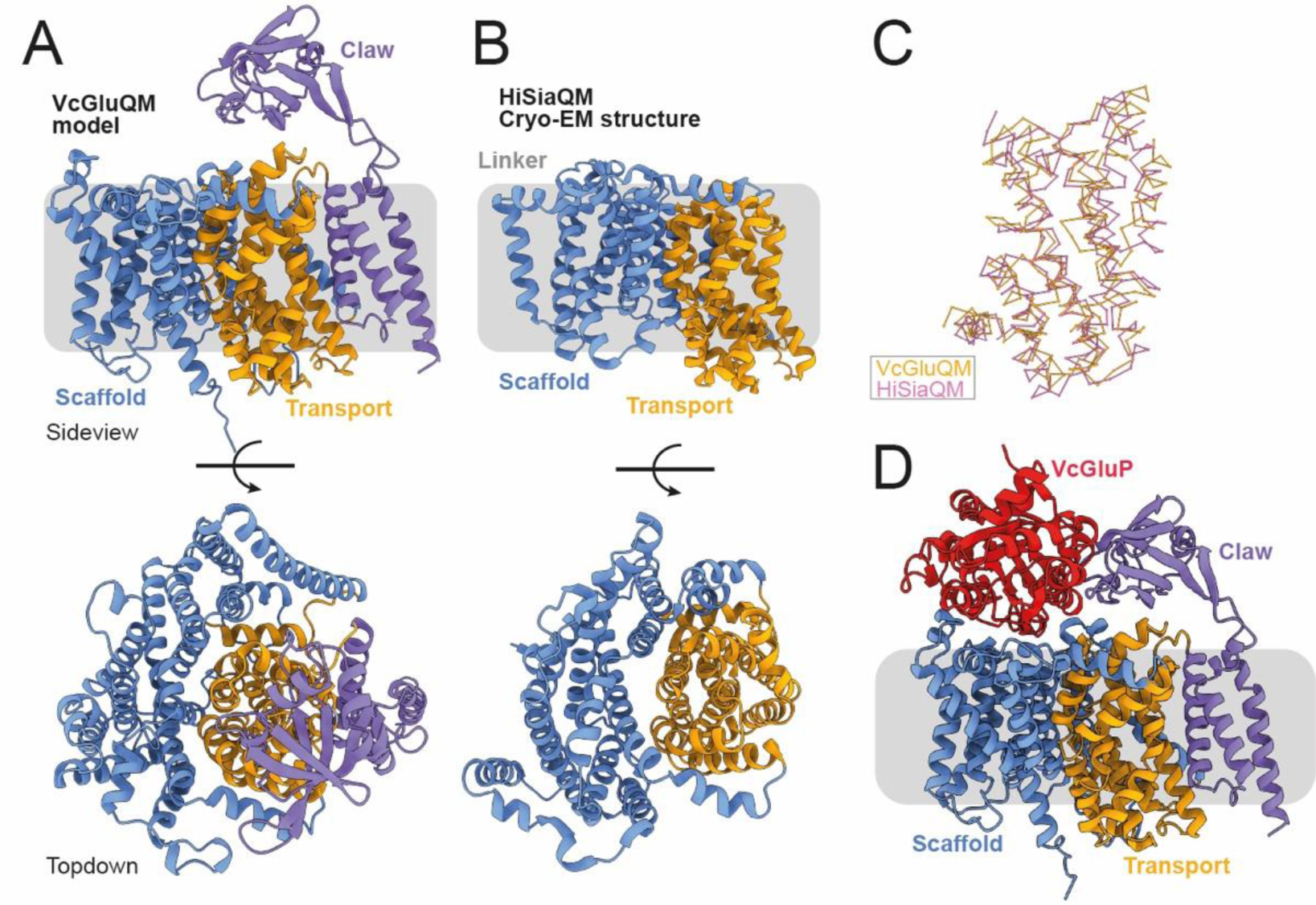
Structural model of VcGluQM and the predicted interactions with VcGluP. **A)** Side view and top-down view of the structural model of VcGluQM generated by Alphafold2. The transport or ‘elevator’ domain is coloured orange, the scaffold is blue, with the light blue indicating the position of the ‘Q’ domain, the linker helix is grey, and the claw domain is colored light purple. **B)** Side view and top-down view of the Cryo-EM structure of HiSiaQM, colored the same as for VcGluQM in A). **C)** Superimposition of the chain traces of the transport domains of VcGluQM (orange) and HiSiaQM (pink). **D)** Cartoon representation and **E)** surface representation of the Colabfold model of the holocomplex VcGluPQM, colored as in A) and with VcGluP colored red.

**Figure 9.**
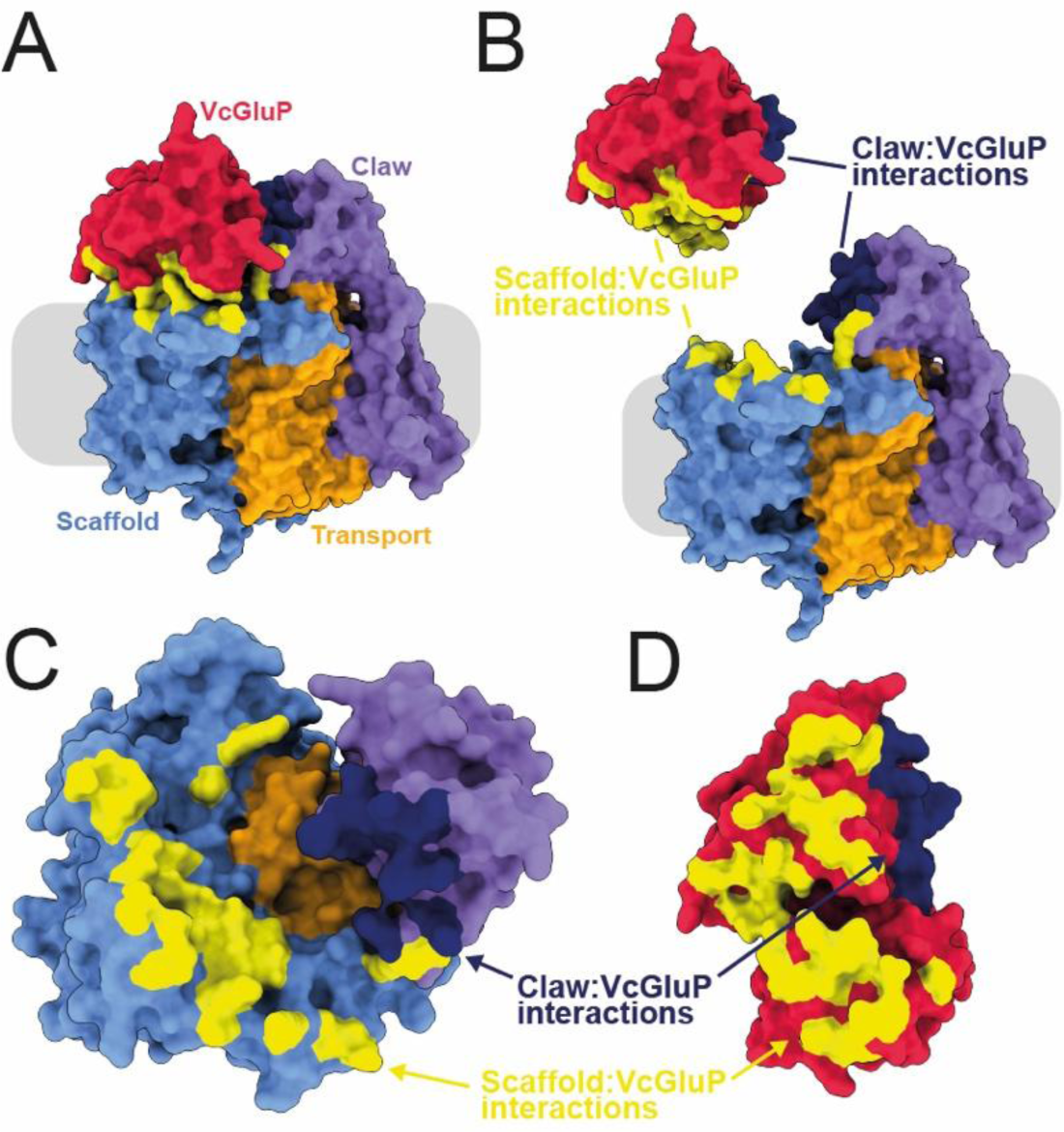
Interfacial interactions between VcGluP and VcGluQM. **A)** Surface representation of VcGluPQM colored as in Figure 8. Interactions between the scaffold domain and VcGluQM are in yellow and the interaction between VcGluP and the claw domain are in dark blue. **B)** Same as A) but interface has been exploded. **C)** Top-down view of VcGluQM showing interfacial contacts, and **D)** bottom-up view of VcGluP showing interfacial contacts.

When modelled independently using Alphafold2, VcGluP adopts a closed state identical to our crystal structure (Supplementary figure 3A). However, superimposition of our crystal structure of the closed, liganded VcGluP and the VcGluP structure predicted during the modelling of the holocomplex with Colabfold reveals that the ligand-free VcGluP in the holocomplex model adopts an open state (Supplementary figure 3B, C and D). With the caveat that these observations are based partially on structural models and require experimental validation, it is tempting to speculate that VcGluP’s open state observed in the Colabfold model is induced by interaction with the membrane component, and may represent a transitional stage of the transport cycle.

## Discussion

In this work, we have structurally characterized a TAXI TRAP SBP from the human pathogen *V. cholerae* and performed in-depth analysis of the substrate binding determinants. We have shown that VcGluP preferentially binds glutamate and can also bind glutamine with a micromolar affinity. Glutamate uptake plays a key role in osmoadaptability of this human pathogen and to the best of our knowledge, this is the only experimentally validated glutamate transporter identified in *V. cholerae*. We have demonstrated that ligand binding involves interactions with side chains and an extensive water network. In addition, we have shown that a glutamine and tyrosine are *the* critical ligand binding residues, and VcGluP lacks a functionally important binding site arginine residue, which is a hallmark of DctP-type TRAP SBPs. Analysis of Alphafold2-based models of the membrane component suggests a unique arrangement hitherto unseen for TRAP transporters, in which a C-terminal extension forms a globular periplasmic domain and makes numerous interactions with the SBP.

This work is the first to investigate the functional contributions of binding site residues in a TAXI TRAP transporter SBP. Characterization of the substrate range and selectivity of DctP-type TRAP transporter SBPs has revealed that they can bind to a diverse range of compounds, the majority of which contain at least one carboxyl group^2,16^. The numerous structures of DctP-type SBPs has revealed that the carboxyl group of the ligand is coordinated by an arginine sidechain, which is highly conserved in DctP-type SBPs and serves as a selectivity filter for binding^26^. Our structure and binding site characterization of VcGluP has revealed that no arginine (or lysine) plays a role in the substrate binding site, revealing a key difference between the two TRAP sub-families.

Binding of L-glutamate by VcGluP is facilitated by a range of sidechain interactions (Fig. 7). In addition, an extensive water network also provides several key interactions (Fig. 7D), which is a feature observed previously in other TRAP SBP binding sites^56^. From our structure, we identified 6 residues potentially involved in ligand binding; Y45, S75, Q93, E125, Y200 and M201. However, we found that binding affinity was only severely diminished when Y45 and Q93 were mutated to alanine, reducing the affinity by ∼150 and ∼750-fold (Table 1), respectively, suggesting these residues are the key binding determinants for VcGluP.

Comparison of VcGluP’s binding site reveals that it is virtually identical to the binding site of TtGluBP (TTHA1157)^37^. While both protein belong to Cluster F^53^, the identical binding site was unexpected considering they did not coalesce on our phylogenetic tree and are only 38% identical to each other (Fig. 1). A feature of both TtGluBP and VcGluP is the positioning of 2 tyrosine residues either side of the glutamate (Y45 and Y200 in VcGluP)^37^. BugE, from the tripartite tricarboxylate transporter (TTT) family of binding protein-dependent secondary active transporters, also binds glutamate, and exhibits the same overall architecture all SBPs share^57^. While there is no sequence similarity between the VcGluP and BugE, and the overall binding site composition is different, both proteins use aromatic residues (Phe in BugE and Tyr in VcGluP) to sandwich the aliphatic chain of the glutamate providing hydrophobic interactions (Fig. 7D)^57^. Aromatic residues are also prominent in the binding site of the pyroglutamate-specific DctP-type TRAP transporters from *Bordetella pertussis*, Dct6 and Dct7, but in these proteins the aromatic residues are a combination of tryptophan and tyrosine^13^. To support the importance of these aromatic residues, the equivalent residue to Y45 is found in 78% of the TAXI sequences in our alignment (Supplementary figure 1). The equivalent residue to Y200 is only found in 20% of sequences reflective of its negligible contribution to binding (Table 1). The residue that appeared to contribute most significantly to binding, Q93, is found in 54% of the sequences in our alignment (Supplementary figure 1), suggesting this is possibly a variable site that will change allow different TAXIs to accommodate different ligands. However, more information on the range of ligands and the binding determinants of TAXI SBPs is required to evaluate this.

The cryo-EM structures of DctP-type TRAP membrane components have revealed important insight into the transport mechanism of TRAP transporters and have confirmed previous predictions^19,24,29^. We currently lack any experimental structures of TAXI-TRAP membrane components. While we show here that they are predicted to largely resemble DctP-type TRAP membrane components, the low sequence identity between DctP- and TAXI-type membrane components will likely result in mechanistic differences. Indeed, significant divergence has also been observed by the TAXI-TRAP from *P. mirabilis* using a H^+^ gradient, unlike the Na^+^-driven DctP-type TRAP transporters that have been characterized^36^. Investigation into the predicted structure of VcGluQM revealed that it is substantially larger than the structurally characterized DctP-type TRAP membrane components (Fig. 8). Most of the size difference between VcGluQM and HiSiaQM is accounted for by VcGluQM’s periplasmic globular C-terminal extension that is predicted to interact with the SBP (Fig. 9). This extension has not been reported for any DctP-type TRAP transporters, suggesting it is a TAXI-specific addition. However, it is highly reminiscent of the similar domain in the maltose ABC transporter, MalEFGK2, but However, the claw domain does not appear to be a general feature of TAXI TRAP membrane components either. Analysis of the Alphafold2-derived models of the membrane components associated with the 59 TAXI transporters in our phylogenetic tree reveals that only the TAXI TRAP transporters in the vicinity of VcGluP are predicted to have this claw domain. All other TAXI transporters in our tree, if they have an associated membrane component gene, lack this claw domain in their membrane components.

The function of the claw domain is currently unknown, but one possibility is that it enhances the affinity between the SBP and membrane component. Recently, analysis of the TAXI family revealed that a large proportion of TAXI SBPs are lipoproteins that are fused via a cysteine to a lipid headgroup^36^. This membrane tethering would substantially increase the local SBP concentration and overcome a low affinity interaction between SBP and membrane component. Analysis of VcGluP’s sequence using SignalP 6.0 reveals that it is predicted with 99.9% confidence to contain a Sec/SPI signal peptide^58^, meaning that VcGluP would move unrestricted in the periplasm rather than being tethered to the membrane. Therefore, in this case, the claw domain may simply be stabilizing the interaction between the membrane component and an untethered SBP. However, further investigation into the spread of the claw domain and how it correlates with membrane tethering is required.

In this work, we have shown that VcGluPQM is likely an L-glutamate transporter from the Gram negative, halophilic intestinal pathogen *Vibrio cholerae* which still poses a significant threat to human life, especially in lower income countries^59^. As *V. cholerae* naturally inhabits estuarine environments, it undergoes substantial fluctuations in the ionic strength of the growth media during its lifecycle, which requires a robust response to avoid cell lysis. While glutamate is itself a known osmolyte^60^, and its uptake by *V. cholerae* increases under elevated osmotic pressure^61^, it is actually used as precursor for ectoine, which is one of the main compatible solutes used by *V. cholerae*^62^. Therefore, glutamate uptake via VcGluPQM likely contributes substantially to osmoadaptability allowing the pathogen to survive in elevated external osmolarity during niche adaptation and in the intestine during infection^62^.

Here, we have characterized VcGluP, a TAXI TRAP SBP from *V. cholerae*, the first experimentally validated glutamate transporter identified in this pathogen. This work has shed new light on the binding determinants of TAXI transporters, provided new insight into the protein:protein interactions of binding protein dependent transporters, and has identified a transporter in a human pathogen that likely influences its environmental adaptability.

## Acknowledgements

We thank Lowri Williams and Rosie Young for technical assistance. This work was financially supported by the BBSRC SoCoBio doctoral training partnership and a BBSRC responsive mode grant (BB/V007424/1) awarded to CM. This work is dedicated to the memory of Bernie Strongitharm.

## Author contributions

NM and JFSD and CM purified the proteins. CK performed the bioinformatic analysis. JFSD collected all binding assay data. AL and BS performed the mass spec analysis. AD and JFSD collect and processed the X-ray data and built the atomic models. AD, JFSD and CM analyzed the structures. MFB, GHT and CM supervised the research. CM wrote the manuscript. All authors read and edited the manuscript.

## Data availability statement

X-ray structures have been deposited in the Protein Data Bank. PDB ID: 8S4J.

## Supplementary information

**Supplementary table 1.**
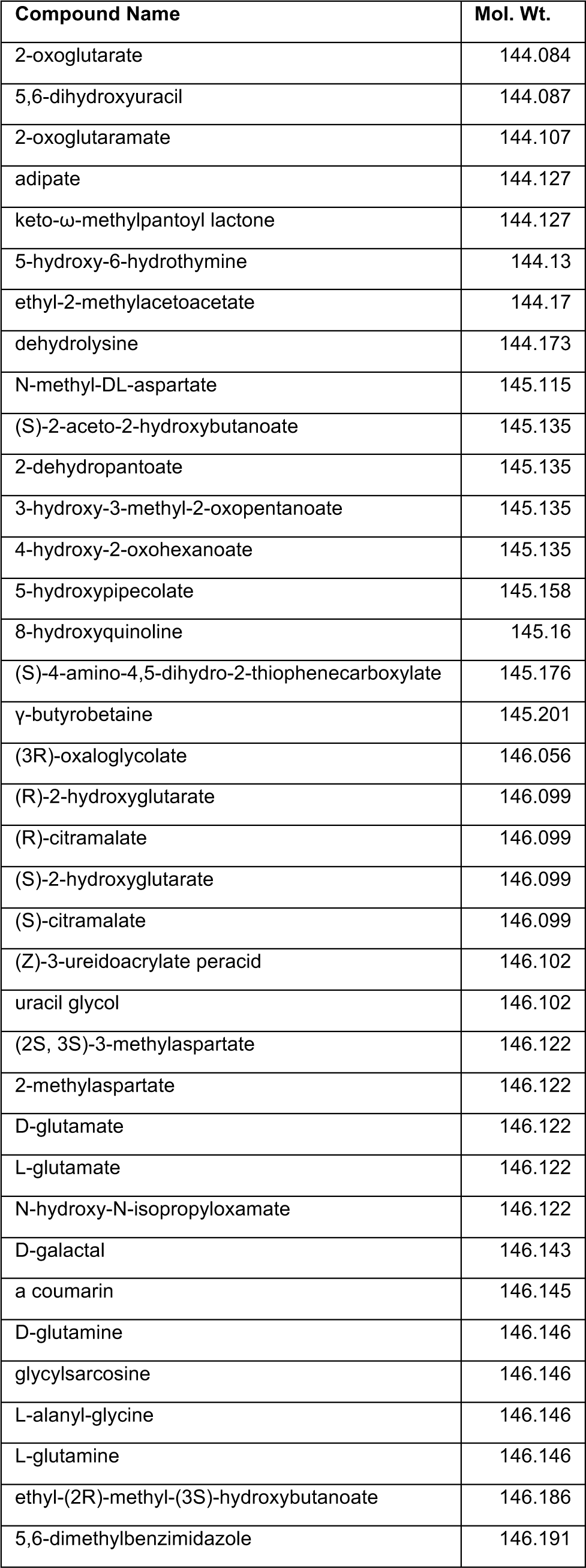
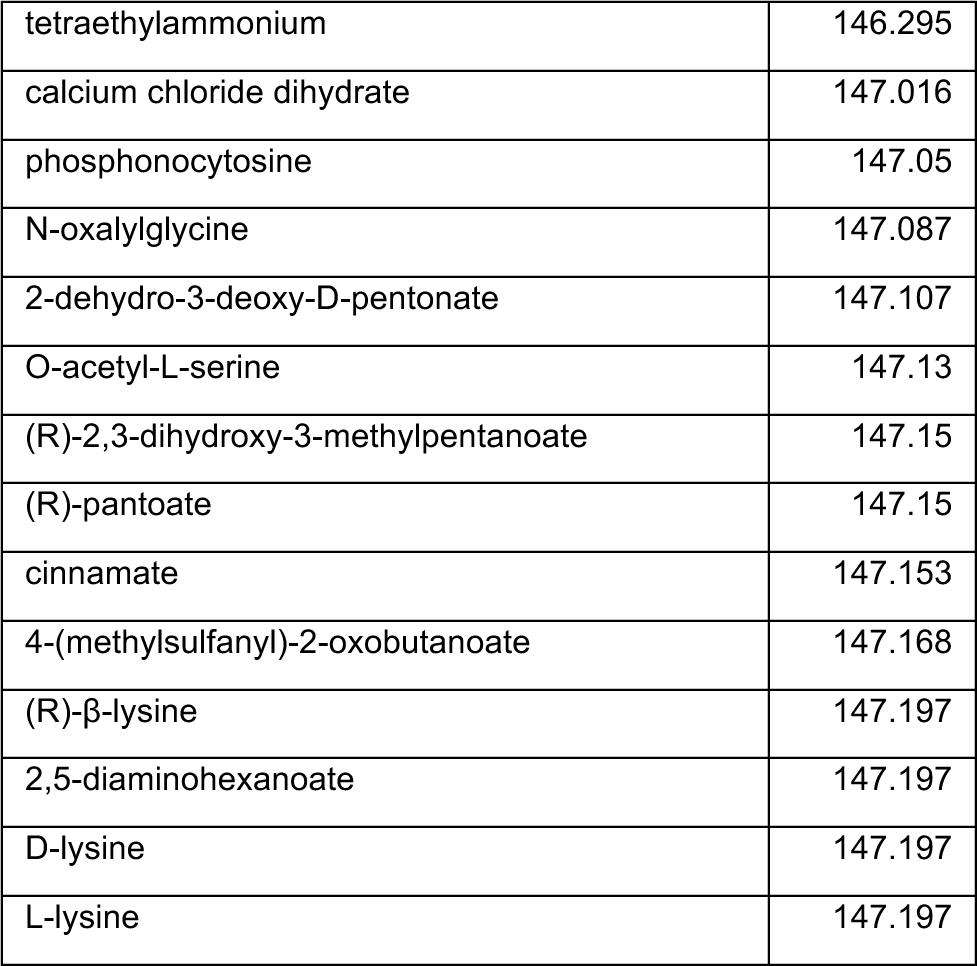
List of E. coli metabolites between 144-148 Da. Data obtained from EcoCyc.org.

**Supplementary Table 2.** Data collection and refinement statistics. Statistics for the highest-resolution shell are shown in parentheses.

|  | **Vc0430 Bound to L-Glutamate** |
| --- | --- |
| **Wavelength** |  |
| **Resolution range** | 69.64 - 1.7 (1.761 - 1.7) |
| **Space group** | R 3 2 :H |
| **Unit cell** | 126.66 126.66 180.28 90 90 120 |
| **Total reflections** | 1242043 (122113) |
| **Unique reflections** | 61144 (6061) |
| **Multiplicity** | 20.3 (20.1) |
| **Completeness (%)** | 99.99 (99.98) |
| **Mean I/sigma (I)** | 26.31 (3.42) |
| **Wilson B-factor** | 27.60 |
| **R-merge** | 0.06415 (0.9057) |
| **R-meas** | 0.0658 (0.9292) |
| **R-pim** | 0.01456 (0.2063) |
| **CC1/2** | 1 (0.917) |
| **CC\*** | 1 (0.978) |
| **Reflections used in refinement** | 61140 (6066) |
| **Reflections used for R-free** | 3035 (317) |
| **R-work** | 0.1902 (0.3618) |
| **R-free** | 0.2136 (0.3516) |
| **CC (work)** | 0.969 (0.824) |
| **CC (free)** | 0.964 (0.809) |
| **Number of non-hydrogen atoms** | 2698 |
| **macromolecules** | 2425 |
| **ligands** | 15 |
| **solvent** | 258 |
| **Protein residues** | 300 |
| **RMS(bonds)** | 0.007 |
| **RMS(angles)** | 0.96 |
| **Ramachandran favored (%)** | 96.97 |
| **Ramachandran allowed (%)** | 3.03 |
| **Ramachandran outliers (%)** | 0.00 |
| **Rotamer outliers (%)** | 1.52 |
| **Clashscore** | 1.65 |
| **Average B-factor** | 28.09 |
| **macromolecules** | 26.92 |
| **ligands** | 27.41 |
| **solvent** | 39.12 |

**Supplementary figure 1.**
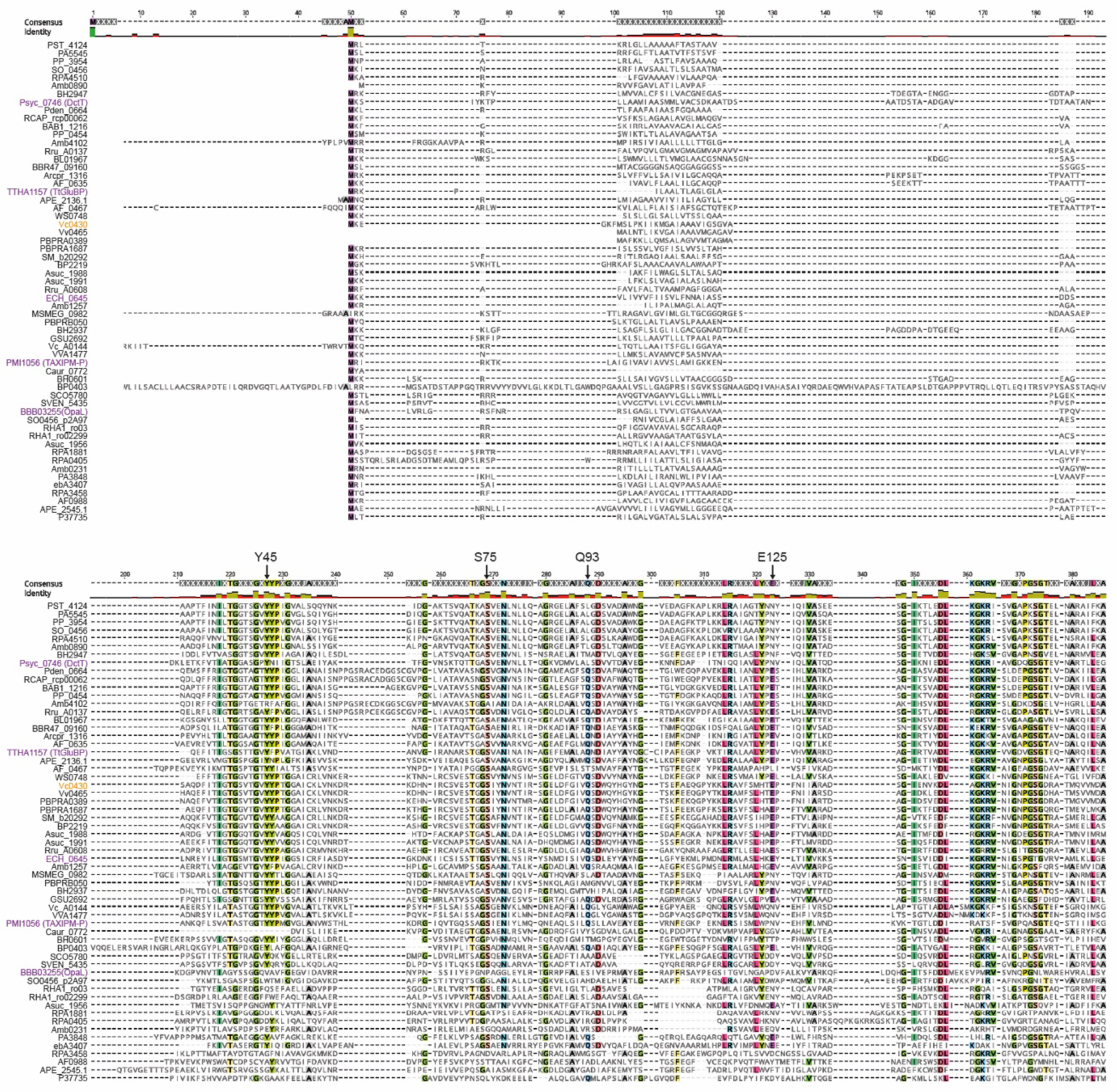
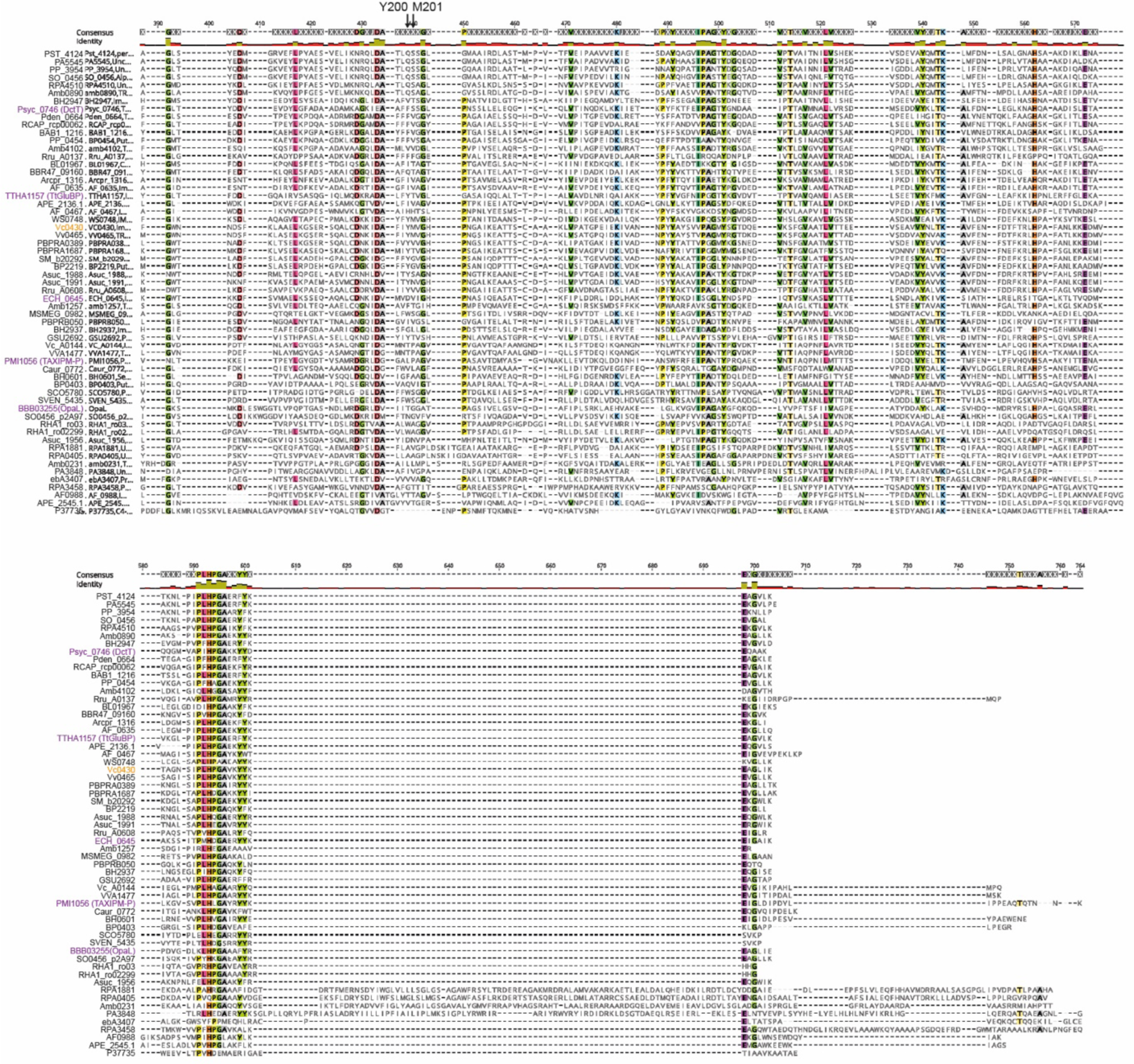
Multiple sequence alignment of 59 TAXI SBPs. Sequence alignment to accompany the cladogram in Figure 1. Protein names are colour coded to match the tree in Figure 1 and the binding site residues mutated to alanine are indicated with an arrow.

**Supplementary figure 2.**
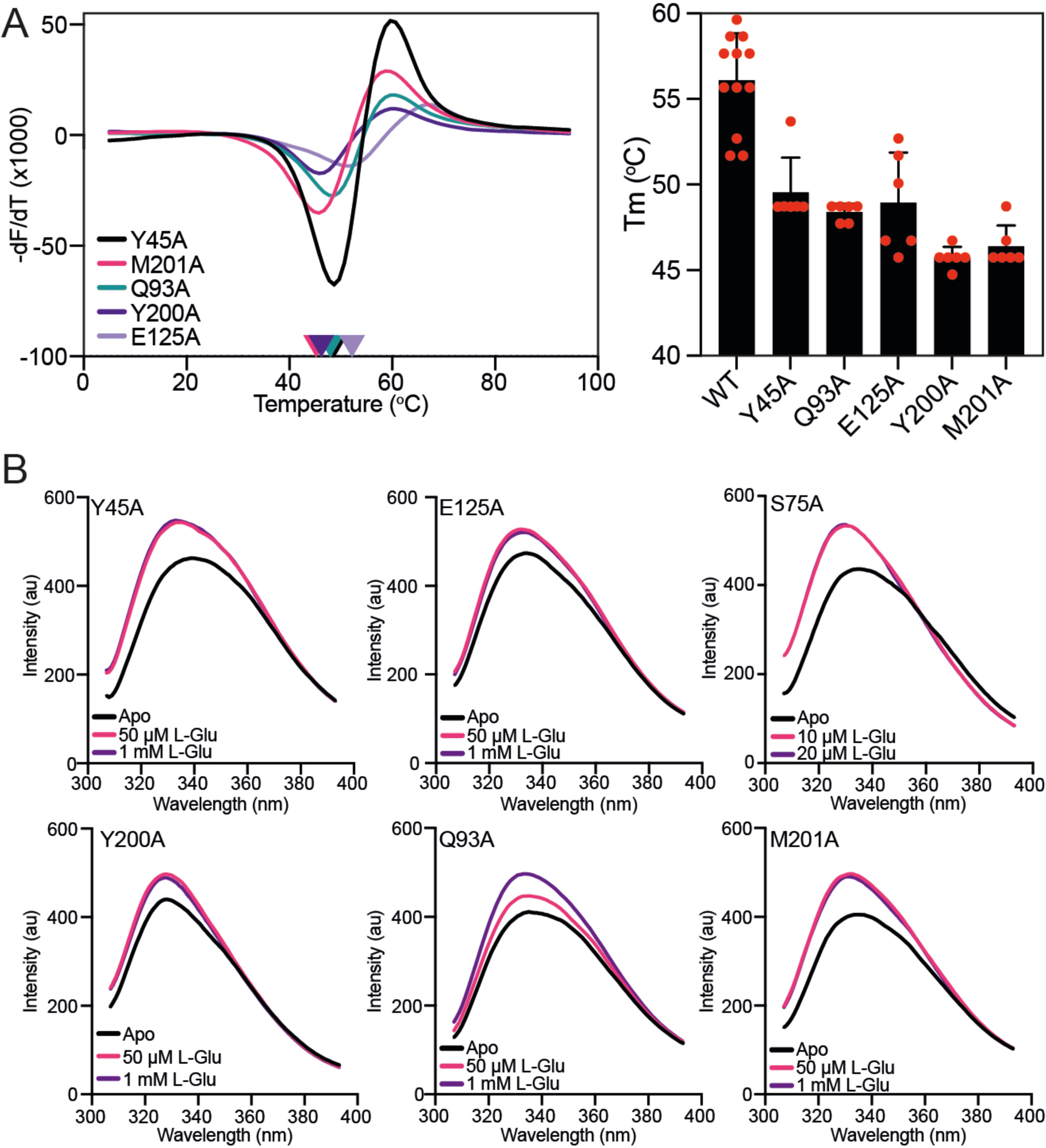
Thermostability and tryptophan fluorescence analysis of VcGluP binding site mutants. **A)** Melting temperature of each VcGluP binding site mutant. Left panel: representative DSF data for each VcGluP binding site mutant in the absence of substrate. Right panel: the Tm of each binding site mutant derived from the DSF assay. Error bars represent standard deviation and individual datapoints are shown in red. **B)** Emission scans of each mutant with excitation wavelength of 295 nm in the presence and absence of L-glutamate.

**Supplementary figure 3.**
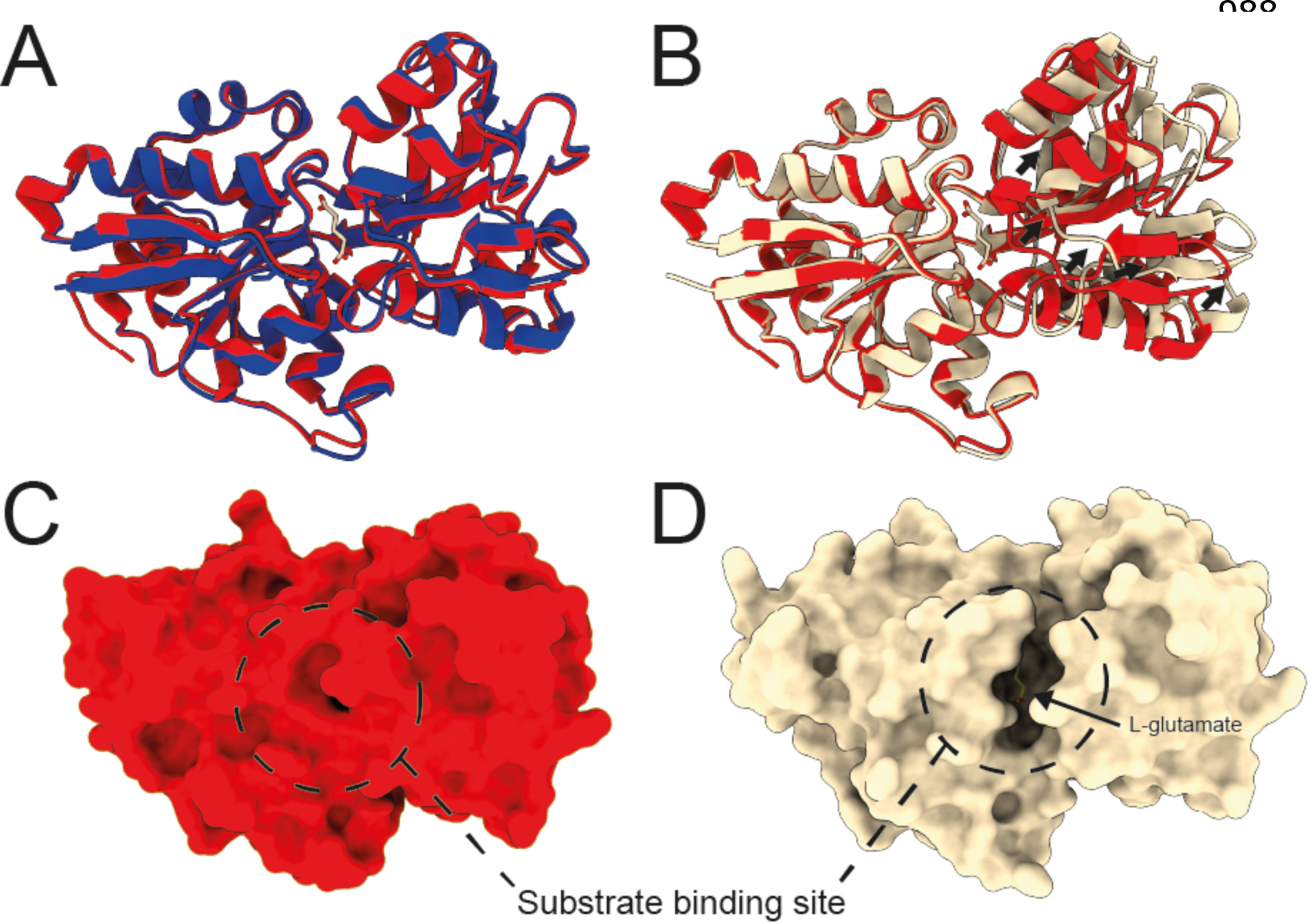
Structural alignments of VcGluP structural models. **A)** Superimposition of the cartoon representations of VcGluP crystal structure (red) or Alphafold2 model (blue). **B)** Superimposition of the cartoon representations of VcGluP crystal structure (red) or model derived from VcGluPQM Colabfold model (wheat). **C)** Surface representation of the VcGluP crystal structure showing no access to binding site. **D)** Surface representation of the VcGluP derived from the VcGluPQM Colabfold model showing a solvent accessible binding site.

